# Covalently linked adenovirus-AAV complexes as a novel platform technology for gene therapy

**DOI:** 10.1101/2024.08.21.609008

**Authors:** Logan Thrasher Collins, Wandy Beatty, Paul Boucher, Buhle Moyo, Michele Alves-Bezerra, Ayrea Hurley, William Lagor, Gang Bao, David T. Curiel, Zhi Hong Lu

## Abstract

Adeno-associated virus (AAV) has found immense success as a delivery system for gene therapy, yet the small 4.7 kb packaging capacity of the AAV sharply limits the scope of its application. In addition, high doses of AAV are frequently required to facilitate therapeutic effects, leading to acute toxicity issues. While dual and triple AAV approaches have been developed to mitigate the packaging capacity problem, these necessitate even higher doses to ensure that co-infection occurs at sufficient frequency. To address these challenges, we herein describe a novel delivery system consisting of adenovirus (Ad) covalently linked to multiple adeno-associated virus capsids as a new way of more efficiently co-infecting cells with lower overall amounts of AAVs. We utilize the DogTag-DogCatcher (DgT-DgC) molecular glue system to construct our AdAAVs and we demonstrate that these hybrid virus complexes achieve enhanced co-transduction of cultured cells, including physiologically relevant primary cells. On this basis, AdAAV technology may eventually facilitate therapeutic co-delivery of multiple transgenes at low virus doses for treating complex ailments.

## Introduction

Although adeno-associated virus (AAV) gene therapy has shown enormous promise and led to eight clinically approved treatments,^1–4^ it has consistently been hampered by the vector’s poor transduction efficiency^5^ as well as by the vector’s low DNA packaging capacity of 4.7 kb.^6^ Relatively inefficient AAV tissue transduction drives the need for high doses in systemic therapies. As a result of these high doses, substantial off-target effects occur, leading to acute toxicity.^5,7^ Dual and triple AAVs have been leveraged to address the packaging capacity issue. In dual and triple AAV systems, a mixture of two or three AAVs carrying distinct DNA cargos are injected with the goal for co-infection of the same cells to occur.^8–13^ But dual and triple AAVs typically require even higher doses than traditional AAV therapies to achieve efficient co-transduction of cells, especially when systemic administration is necessary.^14^ Most dual or triple AAV strategies have therefore focused on applications where local administration into target tissues is possible such as retinal gene therapy.^9,11–13^ As such, it is very difficult to safely utilize AAVs for treating complex diseases which may necessitate multigenic delivery and/or delivery of large split transgenes.^9,11^

While adenovirus (Ad) is thought to have higher immunogenicity than AAV, Ad vectors possess much greater transduction efficiencies compared to AAVs.^15,16^ In this regard, Ad vectors often can be used at lower doses, which may represent a less risky option than high-dose AAV therapies.^17,18^ In addition, Ad immunogenicity can be mitigated by capsid modifications or the utilization of low seroprevalence Ads.^15^ That said, using low doses of Ad can also decrease overall gene expression levels, which is disadvantageous for some types of therapeutics. Ads and AAVs thus have a number of benefits and drawbacks which must be weighed out when designing therapies.

As a platform technology for strong multigenic delivery at low virus doses, we herein constructed an entirely new type of gene delivery system “AdAAV” which physically combines Ad and AAV vectors. This platform consists of a larger (about 100 nm diameter) Ad capsid decorated by numerous smaller (about 25 nm diameter) covalently anchored AAV capsids **(Figure 1A)**. In the wild, adenovirus (Ad) and adeno-associated virus (AAV) are “associated” only by virtue of the AAV’s dependence on Ad machinery to replicate itself.^19^ Their association has not previously involved a physical linkage between the capsids of these types of viruses. By leveraging DogTag-DogCatcher (DgT-DgC) molecular glue peptides,^20^ we have constructed the novel AdAAV complex. In this design, AAVs are covalently anchored to the Ad via an isopeptide bond which spontaneously forms between the DgT and DgC peptides in a highly efficient and specific fashion. We engineered Ad5 to bear a DgC fusion on its pIX capsid protein and we engineered AAV-DJ to possess DgT insertions in all of its VP2 proteins and in some of its VP3 and VP1 proteins. AdAAVs facilitate strong co-transduction of cells **(Figure 1B)** even at low multiplicity of infection (MOI). AdAAVs thereby represent a new vehicle for enhancing cellular co-transduction in AAV gene therapy at low doses.

**Figure 1.**
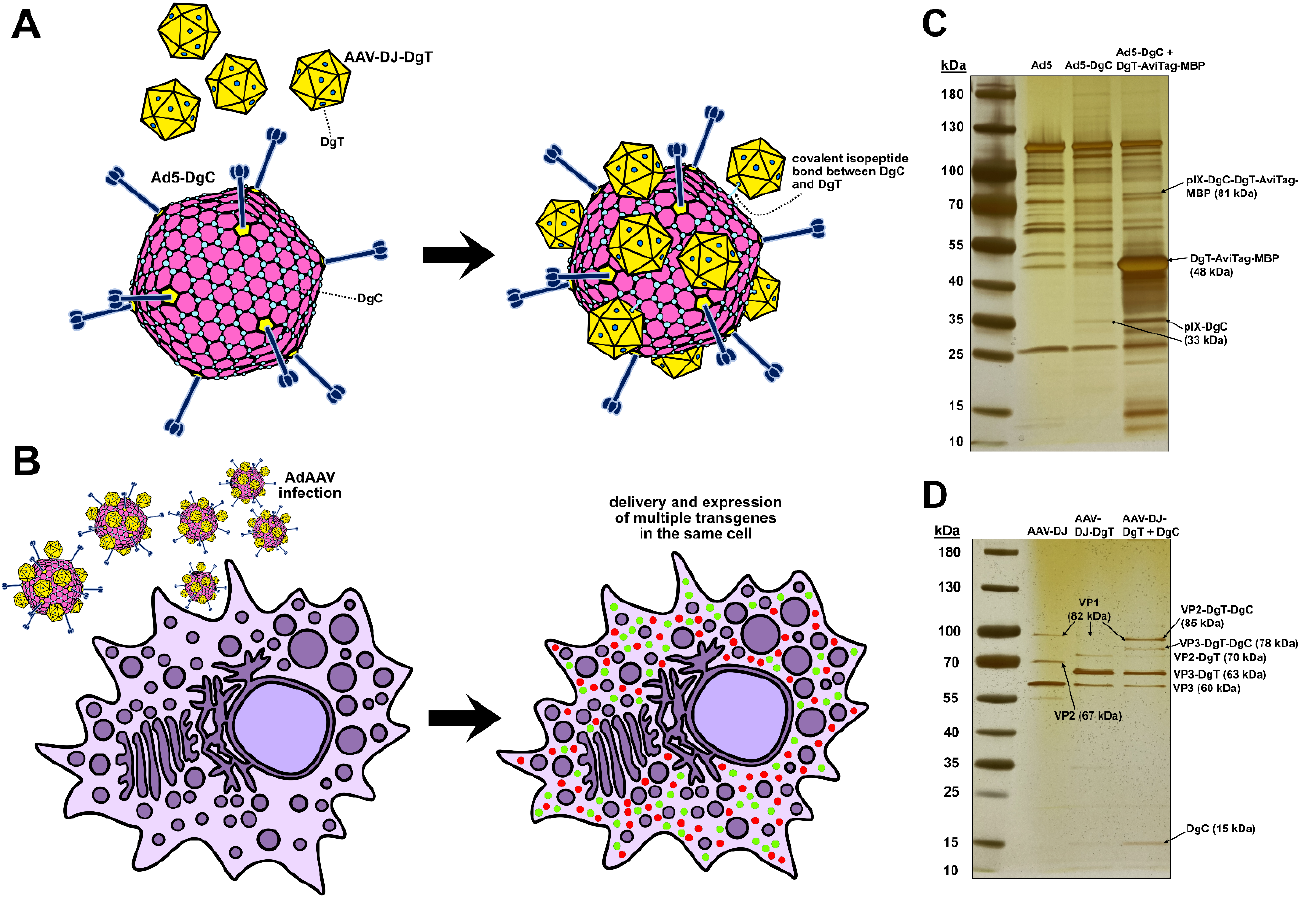
**(A)** Schematic of AdAAV complex assembly and organization. Ad5-DgC has 240 pIX-DgC proteins in its capsid, while AAV-DJ-DgT incorporates a stochastically varying number^26^ of VP2-DgT, VP3-DgT, and VP1-DgT proteins in its capsid. Upon mixing, Ad5-DgC reacts with AAV-DJ-DgT to form covalent isopeptide bonds between their DgC and DgT peptides. **(B)** Schematic of AdAAVs infecting a cell and expressing multiple transgenes as seen in this study. **(C)** Silver stain SDS-PAGE gel of Ad5, Ad5-DgC, and Ad5-DgC mixed with 20-fold excess (relative to pIX-DgC copies) DgT-AviTag-MBP. In the latter lane, the reaction between pIX-DgC and DgT-AviTag-MBP produces a new band at 81 kDa corresponding to pIX-DgC-DgT-AviTag-MBP. **(D)** Silver stain SDS-PAGE gel of AAV-DJ, AAV-DJ-DgT, and AAV-DJ-DgT mixed with 20-fold excess (relative to VP2-DgT copies) DgC. In the latter lane, the reactions of DgC with VP2-DgT and VP3-DgT produce new bands at 78 kDa and 85 kDa. No VP1-DgT or VP1-DgT-DgC is visible, indicating that the proteins were probably very low in copy number.

We envision AdAAV as a way of circumventing the efficiency and dosage limitations of multigenic AAV gene therapy. By physically linking AAVs to an Ad, the platform improves co-transduction efficiency for multiple cargos into the same cell as compared to when the Ads and AAVs are separated. It is important to note that even without physical linkage, the presence of the Ad improves the transduction efficiency of the AAVs, though to a lesser extent. This is likely due to the helper functions provided by the Ad to the AAV. The overall effect is to drastically lower the dose of viruses needed to achieve efficient co-transduction, making AdAAV a promising platform technology for multigenic gene delivery to treat complex diseases.

Though here we only deliver a single AAV cargo and a single Ad cargo to establish proof-of-principle, the high number of AAVs linked to each Ad indicates that the technology should easily extend to co-delivery of both the Ad cargo and more than one AAV cargo. As a platform technology, AdAAV will have immense versatility. In addition to facilitating multigenic gene therapy for complex diseases,^15,21,22^ AdAAVs may enable delivery of multigenic circuits that offer safer and more precise treatments through dynamic responses to changing environmental conditions in real time.^23–25^ Our studies here suggest that AdAAVs will provide a powerful new delivery system to the toolkit of gene therapy with advantages for low-dose multigenic treatments of complex ailments.

## Results

As a first step for constructing AdAAVs, we modified capsid proteins in Ad5 and AAV-DJ to incorporate the DgC and DgT peptides respectively. We decided to fuse DgC to the minor Ad capsid protein pIX since alterations to this protein are typically less disruptive to production and infectivity compared with the other capsid proteins.^27^ With 240 copies of pIX per Ad capsid, the Ad5-DgC virus possesses numerous possible locations for the AAVs to conjugate. We confirmed that the pIX-DgC protein was incorporated into the Ad5-DgC capsid via SDS-PAGE and silver staining **(Fig. 1C)**. In this gel, we also included Ad5-DgC incubated with a 20-fold excess (relative to pIX-DgC copies) of purified DgT-AviTag-MBP protein to verify that the pIX-DgC could react with DgT. By constructing separate pAAV-DJ-VP2-DgT and pAAV-DJ-VP13 plasmids, we inserted DgT into the VP2 protein of AAV-DJ, a structural protein with an average copy number of 5 per capsid. To make sure that only some of the VP3 proteins (average copy number of 50 per capsid) and VP1 proteins (average copy number of 5 per capsid) would bear DgT, we mutated the VP3’s and VP1’s start codons in the pAAV-DJ-VP2-DgT plasmid to the weak start codon CTG. AAVs can utilize CTG^28^ though it leads to less expression compared to ATG. In this way, we constructed an AAV-DJ-DgT vector with VP2-DgT as well as a moderated amount of VP3-DgT, providing a balance between having more available DgT sites and minimizing the negative impact of capsid modifications on production and infectivity. Only a small amount of VP1 was present in the AAV-DJ-DgT capsids (see very faint band), but this did not turn out to preclude infectivity as will be seen later. We could not see VP1-DgT bands, indicating that very little of this protein was produced. We confirmed that AAV-DJ-DgT incorporated VP2-DgT and VP3-DgT using SDS-PAGE with silver staining **(Fig 1D)**. In this gel, we also included AAV-DJ-DgT incubated with a 20-fold excess (relative to VP2-DgT copies) of purified DgC to verify that the VP2-DgT and VP3-DgT could react with DgC. It should be noted that even with the excess of DgC, some unreacted VP2-DgT and VP3-DgT remained, perhaps due to steric shielding effects from the AAV-DJ capsid. To facilitate cell transduction experiments, we constructed our Ad5-DgC to encode eGFP and we provided our AAV-DJ-DgT with a self-complementary genome (to accelerate gene expression) encoding mCherry. We thereby produced and validated the constituent parts of our AdAAV complexes: Ad5-DgC(eGFP) and AAV-DJ-DgT(mCherry).

We next mixed the Ad5-DgC and AAV-DJ-DgT viruses at a 1:10 copy number ratio and incubated overnight at 4°C to induce formation of covalent isopeptide bonds between the viral capsids. Resulting AdAAV complexes were validated using negative stain transmission electron microscopy (TEM) **(Figure 2A)**. We also took TEM images of control mixtures of Ad5 (with unmodified pIX instead of pIX-DgC) and AAV-DJ-DgT at a 1:10 copy number ratio **(Figure 2B)** as well as of Ad5-DgC **(Figure S1A)** alone and AAV-DJ-DgT alone **(Figure S1B)**. AdAAV complexes typically featured around 5-15 discernable attached AAVs per adenovirus across multiple separate experiments. Aggregates of more than one AdAAV could be found occasionally as well **(Figure S2A)**, though these were uncommon. In the Ad5 + AAV-DJ-DgT mixtures, the viruses usually did not contact each other, though occasionally a single AAV-DJ-DgT capsid could be seen adjacent to an Ad5 **(Figure 2B)**. We surmise that this may have occurred due to weak electrostatic association. These data demonstrate that AdAAV complexes can successfully be made by simply mixing the Ad5-DgC and AAV-DJ-DgT viruses.

**Figure 2.**
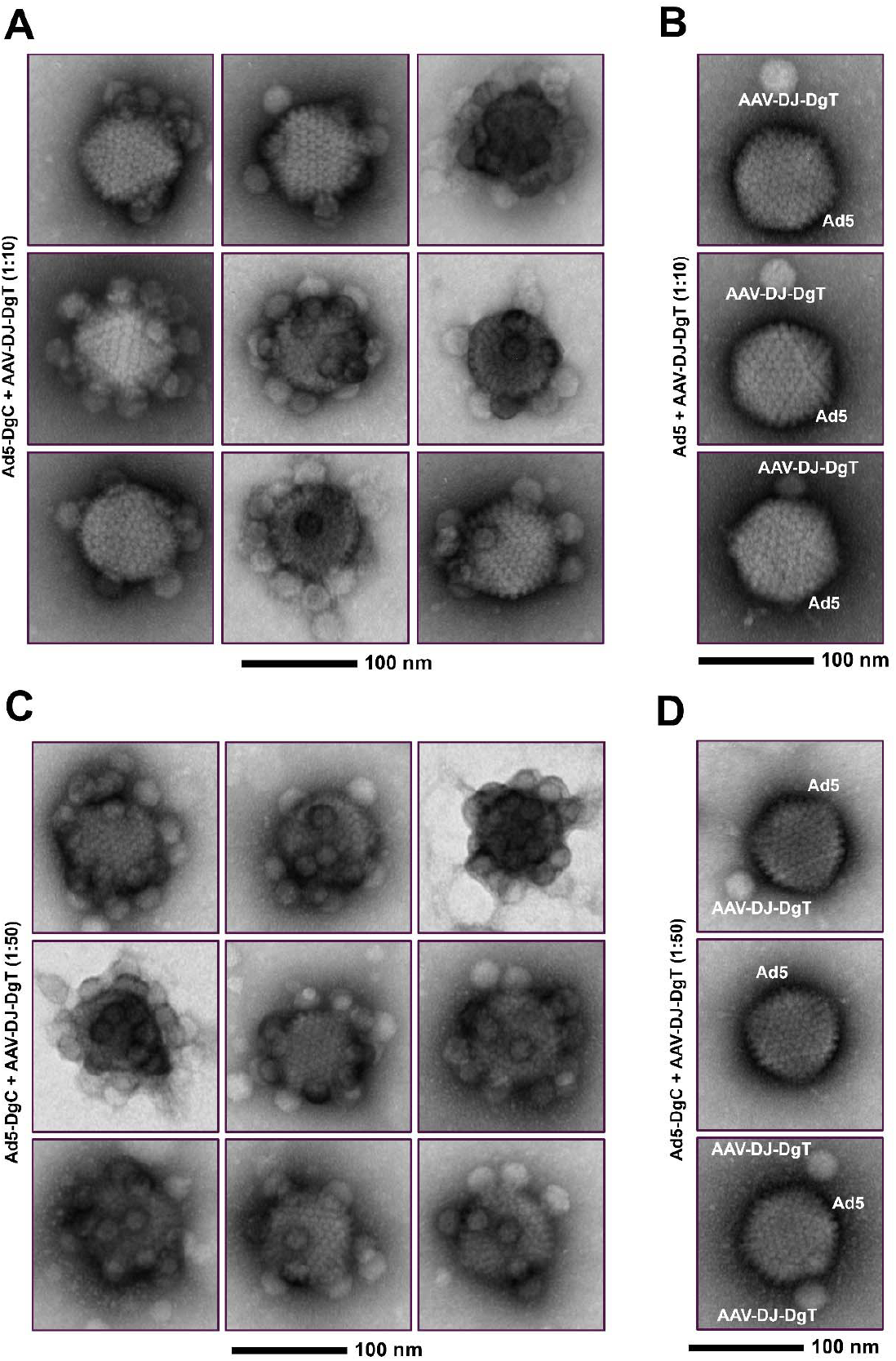
**(A)** AdAAV complexes formed by mixing Ad5-DgC and AAV-DJ-DgT at a 1:10 ratio and incubating overnight at 4°C, visualized using TEM. AdAAVs typically consist of a single Ad5-DgC and 5-15 covalently bound AAV-DJ-DgT. Nine representative AdAAV complexes are shown. Magnification 75,000×. **(B)** Ad5 + AAV-DJ-DgT mixtures subject to the same conditions do not form such complexes. Three representative images of mixtures of Ad5 and AAV-DJ-DgT are shown. Magnification 75,000×. **(C)** AdAAV complexes formed by mixing Ad5-DgC and AAV-DJ-DgT at a 1:50 ratio and incubating overnight at room temperature, visualized using TEM. These AdAAVs typically consist of a single Ad5-DgC and 10-30 covalently bound AAV-DJ-DgT. Nine representative AdAAV complexes are shown. Magnification 75,000×. **(D)** Ad5 + AAV-DJ-DgT mixtures subject to the same conditions do not form complexes. Three representative images of mixtures of Ad5 and AAV-DJ-DgT are shown. Magnification 75,000×.

In an alternative design, we mixed Ad5-DgC and AAV-DJ-DgT viruses at a 1:50 copy number ratio and incubated overnight at room temperature to induce formation of AdAAVs. As before, the resulting complexes were validated by TEM. We observed much more densely saturated AdAAV complexes **(Figure 2C)**. These complexes typically featured around 10-30 discernable attached AAVs per adenovirus across multiple separate experiments. Aggregates of multiple AdAAVs were present but still quite rare **(Figure S2B)**. We also produced control mixtures of Ad5 and AAV-DJ-DgT at a 1:50 copy number ratio. These mixtures again showed vastly less frequent contacts between Ads and AAVs, indicating that the rare colocalizations likely occurred due to chance or weak electrostatic association **(Figure 2D)**. These data illustrate the easy customizability of AdAAV composition through alteration of Ad to AAV ratio.

After constructing our complexes, we tested the capacity of the 1:10 Ad:AAV ratio AdAAVs for co-delivery of two transgenes into the same cell. We infected A549 cells with AdAAV(eGFP, mCherry) complexes, with mixtures of Ad5(eGFP) (bearing unmodified pIX instead of pIX-DgC) and AAV-DJ-DgT(mCherry), with Ad5-DgC(eGFP) alone, and with AAV-DJ-DgT(mCherry) alone. In the AdAAV and Ad5 + AAV-DJ-DgT mixture groups, we used an MOI of 2500 for the Ad component and an MOI of 25000 for the AAV component. In the Ad5 alone group, we used an MOI of 2500. In the AAV-DJ-DgT alone group, we used an MOI of 25000. After 3 days, we imaged the cells via fluorescence microscopy **(Figure S3A)** and utilized flow cytometry to quantify percentages of cells positive for both eGFP and mCherry, for eGFP only, and for mCherry only. For this experiment, the AdAAV group showed successful co-transduction (eGFP+ and mCherry+) across about 48.7% of the cells **(Figure 3A)**. By comparison, the Ad5 + AAV-DJ-DgT group showed co-transduction in about 33.3% of the cells. It is important to realize that the Ad5 virus displayed higher transduction efficiency than the Ad5-DgC virus, so the Ad5 + AAV-DJ-DgT control group’s co-transduction was artificially inflated. Even though this control group had inflated co-transduction, we still observed significantly more enhanced co-transduction with AdAAVs.

**Figure 3.**
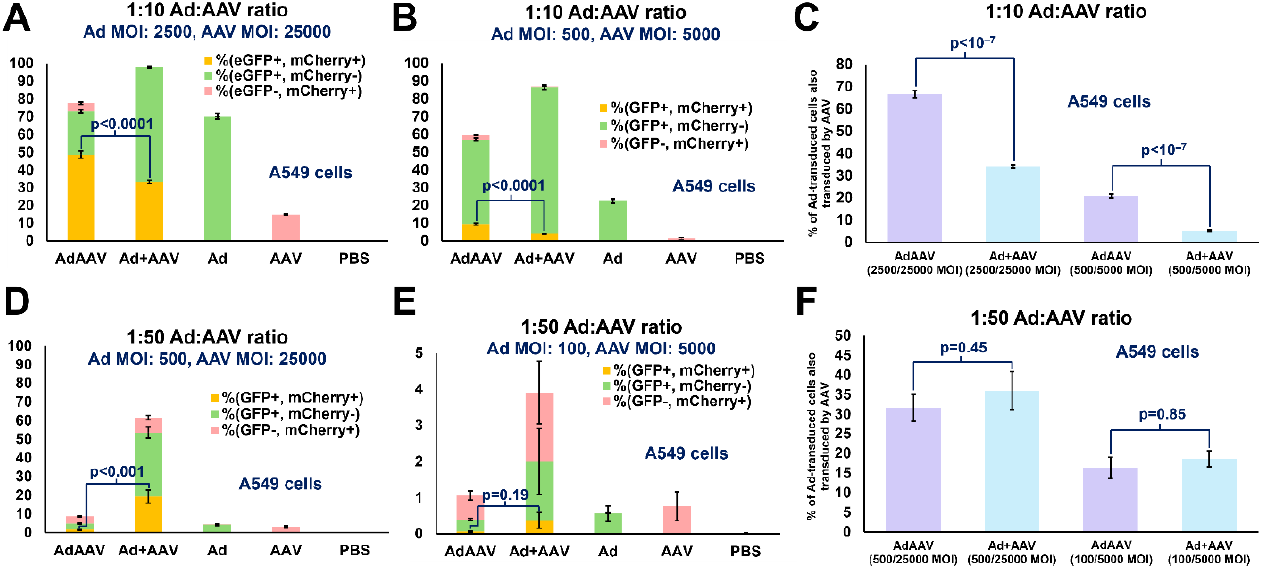
**(A)** Flow cytometry quantification of percentages of A549 cells positive for eGFP and mCherry, for eGFP only, and for mCherry only after infections at 1:10 Ad:AAV ratios. Ad component MOI is 2500 and AAV component MOI is 25000. **(B)** Flow cytometry quantification of percentages of A549 cells positive for eGFP and mCherry, for eGFP only, and for mCherry only after infections at 1:10 Ad:AAV ratios. Ad component MOI is 500 and AAV component MOI is 5000. **(C)** Comparison of the percentages of Ad-transduced cells which were also transduced by AAV across both described MOI levels after infections at 1:10 Ad:AAV ratios. **(D)** Flow cytometry quantification of percentages of A549 cells positive for eGFP and mCherry, for eGFP only, and for mCherry only after infections at 1:50 Ad:AAV ratios. Ad component MOI is 2500 and AAV component MOI is 25000. **(E)** Flow cytometry quantification of percentages of A549 cells positive for eGFP and mCherry, for eGFP only, and for mCherry only after infections at 1:10 Ad:AAV ratios. Ad component MOI is 500 and AAV component MOI is 5000 after infections at 1:50 Ad:AAV ratios. **(F)** Comparison of the percentages of Ad-transduced cells which were also transduced by AAV across both described MOI levels after infections at 1:50 Ad:AAV ratios. For all panels, statistical significance between AdAAV and Ad + AAV groups was evaluated with a two-tailed unpaired t-test. Error bars represent standard error of the mean.

As a test of whether 1:10 Ad:AAV ratio AdAAVs could enhance co-transduction at a lower MOI, we performed the same experiment using fluorescence microscopy **(Figure S3B)** and flow cytometry, but with MOIs of 500 for the Ad components and 5000 for the AAV components. In this case, we saw an even more pronounced difference between the AdAAV group and the Ad5 + AAV-DJ-DgT control. AdAAVs co-transduced about 9.4% of the cells, while Ad5 + AAV-DJ-DgT co-transduced about 4.1% of the cells **(Figure 3B)**. This indicates that AdAAVs may have particular utility for applications where low viral doses are necessary.

To directly examine how well Ad enhanced AAV uptake, we compared the percentages of Ad-transduced cells which were also transduced by AAV in the AdAAV and Ad + AAV groups at both MOIs **(Figure 3C)**. At the 2500/25000 MOI level for AdAAV, 66.6% of cells transduced by Ad were also transduced by AAV. At the 2500/25000 MOI level for Ad + AAV, 34% of cells transduced by Ad were also transduced by AAV. At the 500/5000 MOI level for AdAAV, 20.8% of cells transduced by Ad were also transduced by AAV. At the 500/5000 MOI level for Ad + AAV, 5.3% of cells transduced by Ad were also transduced by AAV. These differences support the idea that AdAAVs have a major advantage in enhancing AAV uptake into cells compared to Ad + AAV, even though Ad + AAV mixtures already improve AAV uptake relative to when AAVs are delivered alone.

We then tested the capacity of the 1:50 Ad:AAV ratio AdAAVs for co-delivery of the eGFP and mCherry transgenes into the same cell, holding constant the AAV MOIs while decreasing the Ad MOIs. We infected A549 cells with AdAAV(eGFP, mCherry) complexes, with mixtures of Ad5(eGFP) (bearing unmodified pIX instead of pIX-DgC) and AAV-DJ-DgT(mCherry), with Ad5-DgC(eGFP) alone, and with AAV-DJ-DgT(mCherry) alone. In the AdAAV and Ad5 + AAV-DJ-DgT mixture groups, we used an MOI of 500 for the Ad component and an MOI of 25000 for the AAV component. In the Ad5 alone group, we used an MOI of 500. In the AAV-DJ-DgT alone group, we used an MOI of 25000. After 3 days, we imaged the cells via fluorescence microscopy **(Figure S4A)** and utilized flow cytometry to quantify percentages of cells positive for both eGFP and mCherry, for eGFP only, and for mCherry only. For this experiment, the AdAAV group showed successful co-transduction (eGFP+ and mCherry+) across only about 1.5% of the cells **(Figure 3D)**. By comparison, the Ad5 + AAV-DJ-DgT group showed co-transduction in about 19.3% of the cells. This demonstrates that oversaturation of Ad capsids with AAVs substantially interferes with AdAAV co-transduction, likely due to steric occlusion of the Ad’s cellular uptake machinery. As such, the 1:10 Ad:AAV ratio is much more desirable than the 1:50 Ad:AAV ratio.

We also tested the 1:50 Ad:AAV ratio AdAAVs could enhance co-transduction at lower MOI. We performed the same experiment using fluorescence microscopy **(Figure S4B)** and flow cytometry, but with MOIs of 100 for the Ad components and 5000 for the AAV components. Just 0.066% of cells were co-transduced in the AdAAV group compared to 0.37% in the Ad + AAV group **(Figure 3E)**. So, the oversaturated AdAAVs again showed decreased co-transduction relative to the Ad + AAV control, providing additional evidence that the 1:10 Ad:AAV ratio is more efficacious than the 1:50 Ad:AAV ratio

We compared the percentages of Ad-transduced cells which were also transduced by AAV in the AdAAV and Ad + AAV groups at both MOIs for the 1:50 Ad:AAV ratio experiments **(Figure 3F)**. At the 500/25000 MOI level for AdAAV, 31.6% of cells transduced by Ad were also transduced by AAV. At the 500/25000 MOI level for Ad + AAV, 36% of cells transduced by Ad were also transduced by AAV. At the 100/5000 MOI level for AdAAV, 16.3% of cells transduced by Ad were also transduced by AAV. At the 100/5000 MOI level for Ad + AAV, 18.6% of cells transduced by Ad were also transduced by AAV. These data further support the idea that the more moderate ratio of 1:10 Ad:AAV is a better strategy.

We next sought to test our AdAAVs in primary human umbilical cord vein endothelial cells (HUVECs), a more challenging target with high physiological relevance. For the 1:10 Ad:AAV ratio, we chose an MOI of 500 for the Ad component of the AdAAV and Ad + AAV groups and an MOI of 5000 for the AAV component of the AdAAV and Ad + AAV groups. In the Ad alone group, we used an MOI of 500. In the AAV alone group, we used an MOI of 5000. After 3 days, we imaged the cells using fluorescence microscopy **(Figure S5A)** and employed flow cytometry to quantify percentages of cells positive for both eGFP and mCherry, for eGFP only, and for mCherry only. The AdAAV group demonstrated significantly higher co-transduction (eGFP+ and mCherry+) across 5.7% of the cells by comparison to the Ad5 + AAV-DJ-DgT group which only co-transduced 1.1% of the cells **(Figure 4A)**. As mentioned earlier, it is important to realize that the Ad5 virus showed higher transduction efficiency than the Ad5-DgC virus, so the Ad5 + AAV-DJ-DgT control group’s co-transduction was artificially inflated. Despite this handicap, we still observed significantly higher co-transduction with AdAAVs. To evaluate how well Ad enhanced AAV uptake, we compared the percentages of Ad-transduced cells which were also transduced by AAV in the AdAAV and Ad + AAV groups. We found for AdAAV that 17.7% of HUVECs transduced by Ad were also transduced by AAV whereas for Ad + AAV only 2% of HUVECs transduced by Ad were also transduced by AAV **(Figure 4B)**. These results in physiologically relevant primary cells support the utility of our AdAAV platform for multigenic delivery.

**Figure 4.**
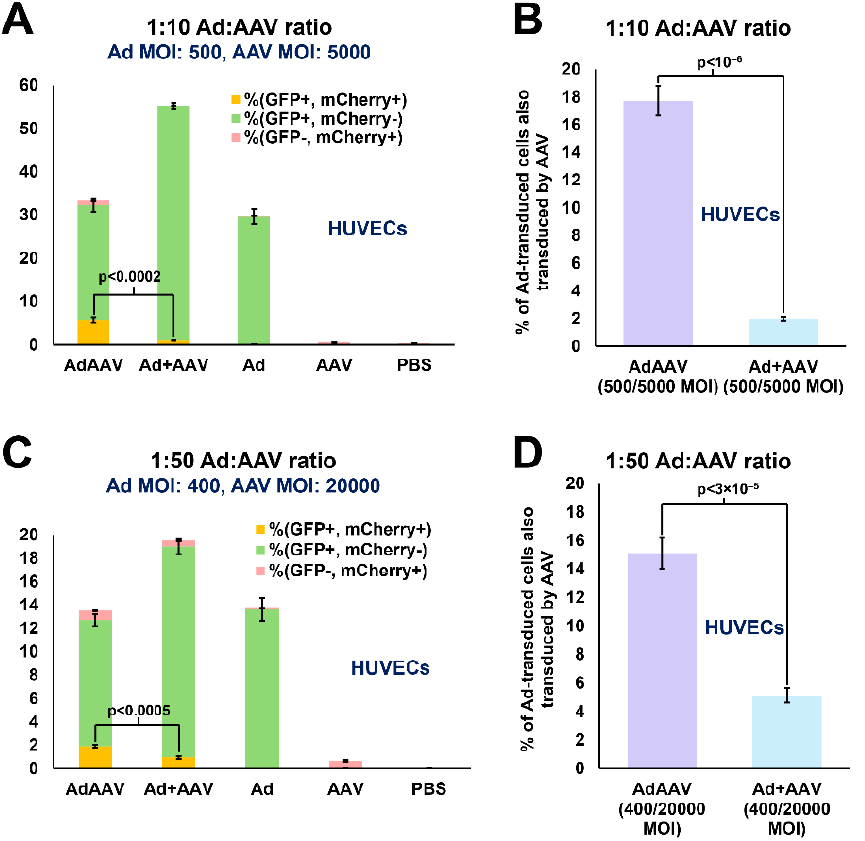
**(A)** Flow cytometry quantification of percentages of HUVECs positive for eGFP and mCherry, for eGFP only, and for mCherry only after infections at 1:10 Ad:AAV ratios. Ad component MOI is 500 and AAV component MOI is 5000. **(B)** Comparison of the percentages of Ad-transduced cells which were also transduced by AAV after infections at 1:10 Ad:AAV ratios. **(C)** Flow cytometry quantification of percentages of HUVECs positive for eGFP and mCherry, for eGFP only, and for mCherry only after infections at 1:50 Ad:AAV ratios. Ad component MOI is 400 and AAV component MOI is 20000. **(D)** Comparison of the percentages of Ad-transduced cells which were also transduced by AAV after infections at 1:50 Ad:AAV ratios. For all panels, statistical significance between AdAAV and Ad + AAV groups was evaluated with a two-tailed unpaired t-test. Error bars represent standard error of the mean.

We also tested the 1:50 Ad:AAV ratio batch in HUVECs to determine if the effects of Ad capsid oversaturation with AAVs would persist across cell types. We chose an MOI of 400 for the Ad component of the AdAAV and Ad + AAV groups and an MOI of 20000 for the AAV component of the AdAAV and Ad + AAV groups. In the Ad alone group, we used an MOI of 400. In the AAV alone group, we used an MOI of 20000. After 3 days, we imaged the cells using fluorescence microscopy **(Figure S5B)** and employed flow cytometry to quantify percentages of cells positive for both eGFP and mCherry, for eGFP only, and for mCherry only. The AdAAV group demonstrated significantly higher co-transduction (eGFP+ and mCherry+) across 1.9% of the cells by comparison to the Ad5 + AAV-DJ-DgT group which only co-transduced 1% of the cells **(Figure 4C)**. Although this degree of co-transduction enhancement is weaker than that seen in the 1:10 Ad:AAV ratio case, it is noteworthy that enhancement did occur in HUVECs, unlike in the 1:50 Ad:AAV ratio case for A549 cells. To evaluate how well Ad enhanced AAV uptake, we compared the percentages of Ad-transduced cells which were also transduced by AAV in the AdAAV and Ad + AAV groups. We found for AdAAV that 15.1% of HUVECs transduced by Ad were also transduced by AAV whereas for Ad + AAV 5.1% of HUVECs transduced by Ad were also transduced by AAV **(Figure 4B)**. These results demonstrate that 1:10 Ad:AAV ratio AdAAV construction is still more favorable compared to 1:50 ratio AdAAVs in the context of HUVECs.

Finally, we wanted to demonstrate directly the capacity for the Ad transduction apparatus within AdAAVs to drive the uptake of AAVs into cells that would otherwise be less permissive for AAV transduction. We hypothesized that Ad transduction apparatus plays a central role in driving uptake of AdAAV complexes into cells and in “dragging the AAVs along for the ride”. To test this, we utilized CHO-CAR cells (expressing the CAR receptor) and CHO cells. CAR is the Coxsackievirus and Adenovirus Receptor, a protein which binds Ad5 fiber knob and facilitates Ad transduction.^29,30^ By contrast, CAR is not known to facilitate AAV transduction.^31^ Since we earlier showed the advantages of 1:10 Ad:AAV ratio AdAAVs, we decided to focus on the 1:10 ratio AdAAVs for this experiment. We chose an MOI of 1000 for the Ad component of the AdAAV and Ad + AAV groups and an MOI of 10000 for the AAV component of the AdAAV and Ad + AAV groups. In the Ad alone group, we used an MOI of 1000. In the AAV alone group, we used an MOI of 10000. After 3 days, we imaged the CHO-CAR cells and CHO cells **(Figure S6A-B)** using fluorescence microscopy and employed flow cytometry to quantify percentages of CHO-CAR cells and CHO cells **(Figure 5A-D)** positive for both eGFP and mCherry, for eGFP only, and for mCherry only.

**Figure 5.**
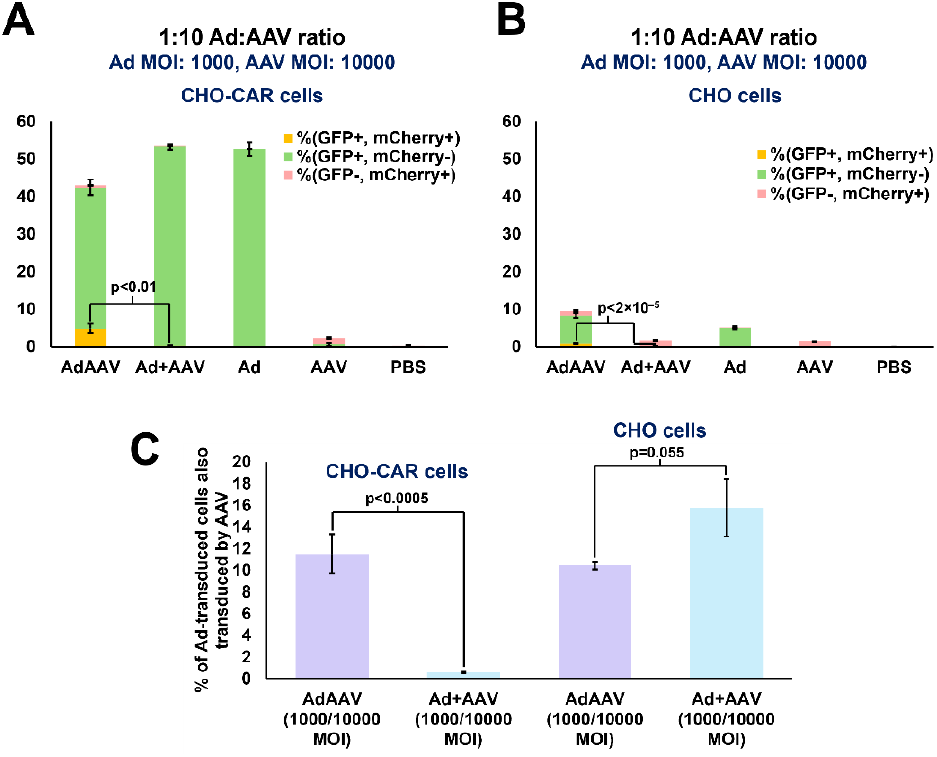
**(A)** Flow cytometry quantification of percentages of CHO-CAR cells positive for eGFP and mCherry, for eGFP only, and for mCherry only after infections at 1:10 Ad:AAV ratios. Ad component MOI is 1000 and AAV component MOI is 10000. **(B)** Flow cytometry quantification of percentages of CHO cells positive for eGFP and mCherry, for eGFP only, and for mCherry only after infections at 1:10 Ad:AAV ratios. Ad component MOI is 1000 and AAV component MOI is 10000. **(C)** Comparison of the percentages of Ad-transduced CHO-CAR cells and CHO cells which were also transduced by AAV after infections at 1:10 Ad:AAV ratios. For all panels, statistical significance between AdAAV and Ad + AAV groups was evaluated with a two-tailed unpaired t-test. Error bars represent standard error of the mean.

For the CHO-CAR cells, the AdAAV group demonstrated significantly higher co-transduction (eGFP+ and mCherry+) across 4.9% of the cells by comparison to the Ad5 + AAV-DJ-DgT group which only co-transduced 0.3% of the cells **(Figure 5A)**. We also once again calculated the percentages of Ad-transduced cells which were also transduced by AAV in the AdAAV and Ad + AAV groups. We found for AdAAV that 11.5% of CHO-CAR cells transduced by Ad were also transduced by AAV whereas for Ad + AAV only 0.6% of CHO-CAR transduced by Ad were also transduced by AAV **(Figure 5C)**. These data strongly indicate that the Ad machinery of AdAAVs drives transduction in the context of Ad-targeted cells which do not otherwise efficiently take up AAVs.

For the CHO cells, very little overall transduction was observed as expected. Although the AdAAV group did demonstrate significantly higher co-transduction (eGFP+ and mCherry+) across 0.86% of the cells in comparison to the Ad5 + AAV-DJ-DgT group which co-transduced 0.03% of the cells **(Figure 5B)**, the very low percentages support the idea that the Ad’s machinery is essential for driving efficient AdAAV cellular transduction. We then calculated the percentages of Ad-transduced cells which were also transduced by AAV in the AdAAV and Ad + AAV groups. We found for AdAAV that 10.5% of CHO cells transduced by Ad were also transduced by AAV whereas for Ad + AAV 15.8% of CHO transduced by Ad were also transduced by AAV **(Figure 5D)**. This result indicates that the lack of an Ad receptor prevented the Ad within AdAAVs from driving uptake of either virus into the cell. These data provide additional evidence that the Ad machinery of AdAAVs drives transduction in the context of Ad-targeted cells which do not otherwise take up AAVs efficiently.

## Discussion

Our data suggest that AdAAV can act as a new platform technology for gene therapy, particularly in the area of enhancing co-delivery of split genetic cargos. Our design represents a next-generation class of hybrid viral vector which goes beyond past genomic mosaic strategies^32^ by instead leveraging physical linkage of distinct capsids. We demonstrate incorporation of pIX-DgC into the Ad5 capsid and VP2-DgT and VP3-DgT into the AAV-DJ capsid via SDS-PAGE silver stain. We also utilize SDS-PAGE silver stain to show that the pIX-DgC reacts to form a covalent linkage with purified DgT-AviTag-MBP as well as that VP2-DgT and VP3-DgT react to form covalent linkages with purified DgC. We validate successful formation of AdAAV complexes by TEM. We then demonstrate enhanced co-delivery of two transgenes, eGFP in the Ad component and mCherry in the AAV component. We showed enhanced co-delivery results in immortalized A549 tumor cells and primary HUVECs, indicating broad physiological relevance for AdAAV constructs. We show that AdAAVs made using a 1:10 Ad:AAV ratio achieve much better results than those made using a 1:50 Ad:AAV ratio, likely due to the greater occlusion of adenoviral transduction machinery in the latter case. We additionally show via comparing infection of CHO-CAR and CHO that the Ad component’s targeting and transduction machinery drives uptake into cells which are not otherwise permissive for AAVs, a finding which suggests that future *in vivo* AdAAV efforts might take advantage of the Ad’s high targetability.^33^ Because each AdAAV consists of an Ad decorated by many AAVs, AdAAVs likely could deliver a mixture of several distinct AAV cargos in addition to Ad cargo. AdAAVs might also eventually deliver split gene products in a fashion similar to that of dual and triple AAVs, but with the advantage of greatly enhanced co-delivery efficiency.

As a future direction, we envision AdAAVs as a superlative platform for multigenic CRISPR-Cas gene insertion. Because of the double-stranded DNA (dsDNA) genome of Ad, any Ad vector on its own will have difficulty with efficient CRISPR-mediated knock-in of genes via homology directed repair (HDR).^34^ Indeed, dsDNA templates are less efficient and more prone to off-target integration. However, Ad’s higher packaging capacity makes it better suited for carrying large Cas protein genes in addition to several guide RNA (gRNA) genes. By contrast, AAV features a single-stranded DNA (ssDNA) genome with a lower packaging capacity. Further iterations of AdAAV may therefore capitalize on the advantages of both vectors, leveraging the Ad to encode a Cas protein and several gRNAs while employing a mixture of AAVs to carry multiple ssDNA donor templates. Alternatively, the Ad component could encode base editors, prime editors, epigenome editors, recombinases, or other machinery that are too large to fit into a single AAV capsid.^35^ CRISPR AdAAVs may someday programmably and efficiently insert multiple transgenes into targeted locations within the human genome, facilitating powerful gene editing therapies for complex diseases.

## Methods

### Production of Ad5-DgC

We made Ad5-DgC by first cloning a fragment containing a linker-pIX-DgC sequence directly into an Ad5 genomic plasmid (E1/E3-deleted Ad5 backbone) via Gibson assembly. The linker was a 45 amino acid rigid spacer derived from the linkers described by Vellinga et al.^36^ After verifying sequence correctness of the resulting Ad5-DgC genomic DNA, we linearized the plasmid and transfected it into HEK293 cells. Serial passages of infection with cell lysate supernatant (lysis was achieved by 4 freeze-thaw cycles per passage) were used to upscale the virus into larger and larger cell culture vessels. We eventually infected 20 T175 flasks with the lysate supernatant from a single T175 flask. After observing approximately 50% cell detachment, we pelleted the cells, performed a final set of 4 freeze-thaw cycles, and loaded the supernatant onto CsCl gradients consisting of 4 mL layer of 1.33 g/cm^3^ CsCl on top of a 4 mL layer of 1.45 g/cm^3^ CsCl. We ultracentrifuged the gradient at 20,000 rpm for 4 hours and then extracted the band corresponding to intact Ad5-DgC particles. This band was loaded onto another CsCl gradient of the same composition and ultracentrifuged at 25,000 rpm for 18 hours. The band was extracted and injected into a Thermo Scientific Slide-A-Lyzer G2 Dialysis Cassette. Dialysis was performed in 1 L of PBS with 10% glycerol buffer for 2 hours before exchanging for fresh buffer to facilitate another 2 hours of dialysis. Finally, the purified Ad5-DgC was withdrawn from the dialysis cassette, aliquoted into 1.5 mL tubes, and stored at –80°C until use. Titration of Ad5-DgC was accomplished using UV-visible spectroscopy.

### Production of AAV-DJ-DgT

To make AAV-DJ-DgT, we first constructed pAAV-DJ-VP13 and pAAV-DJ-VP2-DgT plasmids through cloning with Gibson assembly. As part of our cloning strategy and to ensure that only a moderate fraction of the VP3 and VP1 proteins would be expressed as VP3-DgT and VP1-DgT, we mutated the VP3 and VP1 start codons from ATG to CTG in the pAAV-DJ-VP2-DgT plasmid. For the pAAV-DJ-VP13 plasmid, we used a mutation in the VP2 start codon from ACG to GCG to ensure that expression of only VP1 and VP3 would occur. For the pAAV-DJ-VP2-DgT plasmid, we added a sequence encoding a pair of short flexible linkers flanking the DgT peptide after position 453 in the *Cap* sequence. We employed a self-complementary transfer plasmid encoding mCherry driven by a CAG promoter, pscAAV(mCherry). Next, AAV-DJ-DgT was produced with the help of the viral vector core at UMass Chan Medical School. The core facility transfected HEK293T cells with 0.3 mg of pAAV-DJ-VP13, 1.5 mg of pAAV-DJ-VP2-DgT, 0.75 mg of pscAAV(mCherry), and 0.75 mg of a pAdΔF6 helper plasmid before purification by iodixanol gradient ultracentrifugation. Titration of the self-complementary AAV-DJ-DgT was accomplished using digital droplet PCR.

### SDS-PAGE silver stain, AdAAV assembly, and TEM validation

SDS-PAGE was carried out by running samples on Criterion™ precast gels from Bio-Rad at 50 mA for 1 hour. After removing the gels from their holders, we performed silver stains using a Pierce™ silver stain kit. The first batch of AdAAVs was assembled by mixing Ad5-DgC(eGFP) with self-complementary AAV-DJ-DgT(mCherry) at a 1:10 ratio and incubating overnight at 4°C. Ad5(eGFP) + self-complementary AAV-DJ-DgT(mCherry) controls as well as Ad5-DgC(eGFP) only controls and self-complementary AAV-DJ-DgT(mCherry) only controls were mixed and incubated in the same way. The second batch of AdAAVs was assembled by mixing Ad5-DgC(eGFP) with self-complementary AAV-DJ-DgT(mCherry) at a 1:50 ratio and incubating overnight at room temperature. Ad5(eGFP) + self-complementary AAV-DJ-DgT(mCherry) controls as well as Ad5-DgC(eGFP) only controls and self-complementary AAV-DJ-DgT(mCherry) only controls were mixed and incubated in the same way. For transmission electron microscopy (TEM) validation, a small volume of each group (Ad5-DgC + AAV-DJ-DgT, Ad5 + AAV-DJ-DgT, Ad5-DgC only, AAV-DJ-DgT only) was transferred to a separate tube, diluted 1:10 in PBS with 10% glycerol, negatively stained using 1% uranyl acetate, and imaged by TEM.

### Co-transduction assays

To measure the efficacy of co-transduction in A549 cells (ATCC CCL-185), 1×10^4^ cells per well were seeded in a 96-well plate 4 hours before infection. AdAAV, Ad5 + AAV-DJ-DgT (Ad +AAV), Ad5-DgC only, and AAV-DJ-DgT only were each added to five replicate wells containing cells at the MOI values specified in the Results section. 3 days after infection, we imaged the cells by fluorescence microscopy. Next, the cells were detached using trypsinization and flow cytometry was utilized to quantify the percentages in each well of cells positive for both eGFP and mCherry, of cells positive for only eGFP, of cells positive for only mCherry, and of cells positive for neither. Statistical significance of co-transduction comparisons between AdAAV and Ad5 + AAV-DJ-DgT (Ad + AAV) groups was evaluated using a two-tailed unpaired t-test.

To measure the efficacy of co-transduction in HUVECs (ATCC PCS-100-010), 2×10^4^ cells per well were seeded in a 96-well plate 4 hours before infection. AdAAV, Ad5 + AAV-DJ-DgT (Ad +AAV), Ad5-DgC only, and AAV-DJ-DgT only were each added to five replicate wells containing cells at the MOI values specified in the Results section. 3 days after infection, we imaged the cells by fluorescence microscopy. Next, the cells were detached using trypsinization and flow cytometry was utilized to quantify the percentages in each well of cells positive for both eGFP and mCherry, of cells positive for only eGFP, of cells positive for only mCherry, and of cells positive for neither. Statistical significance of co-transduction comparisons between AdAAV and Ad5 + AAV-DJ-DgT (Ad + AAV) groups was evaluated using a two-tailed unpaired t-test.

To measure the efficacy of co-transduction in CHO-CAR cells and CHO cells (a gift from the laboratory of Anja Ehrhardt), 1×10^4^ cells per well were seeded in a 96-well plate 4 hours before infection. AdAAV, Ad5 + AAV-DJ-DgT (Ad +AAV), Ad5-DgC only, and AAV-DJ-DgT only were each added to five replicate wells containing cells at the MOI values specified in the Results section. 3 days after infection, we imaged the cells by fluorescence microscopy. Next, the cells were detached using trypsinization and flow cytometry was utilized to quantify the percentages in each well of cells positive for both eGFP and mCherry, of cells positive for only eGFP, of cells positive for only mCherry, and of cells positive for neither. Statistical significance of co-transduction comparisons between AdAAV and Ad5 + AAV-DJ-DgT (Ad + AAV) groups was evaluated using a two-tailed unpaired t-test.

## Acknowledgements

We thank the viral vector core at UMass Chan Medical School for their assistance in producing AAV-DJ-DgT vectors. We thank Professor Anja Ehrhardt’s laboratory for their generous gift of CHO-CAR cells and CHO cells.

