## Supplemental Information for "Covalently linked adenovirus-AAV complexes as a novel platform technology for gene therapy"

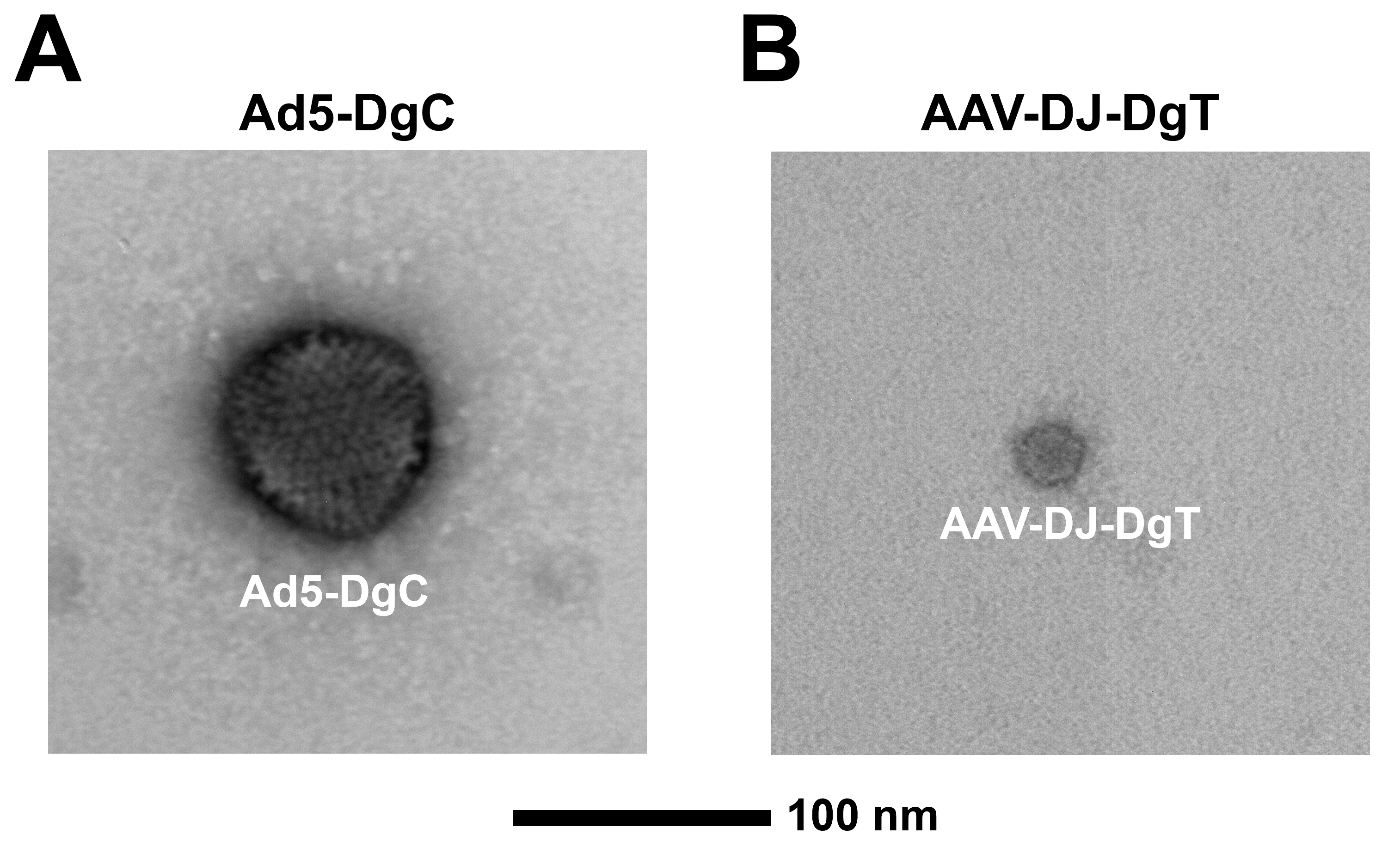

**Supplemental Figure 1 (A)** Negative stain transmission electron microscopy (TEM) image of an Ad5-DgC virus on its own. **(B)** Negative stain TEM image of an AAV-DJ-DgT virus on its own.

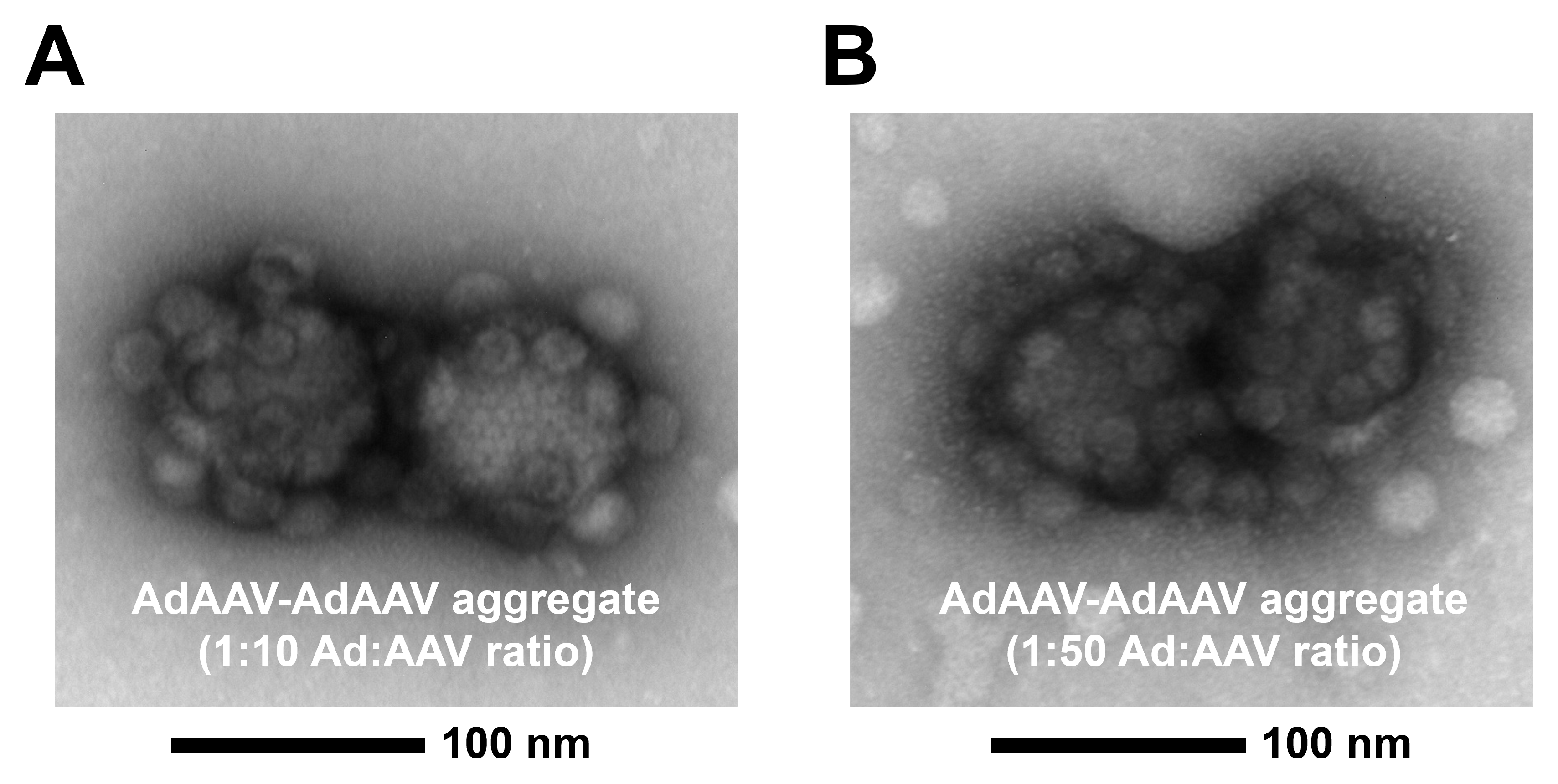

**Supplemental Figure 2 (A)** Negative stain TEM image of a relatively rare aggregate consisting of two AdAAVs after mixing Ad5-DgC and AAV-DJ-DgT at an Ad:AAV ratio of 1:10. **(A)** Negative stain TEM image of a relatively rare aggregate consisting of two AdAAVs after mixing Ad5-DgC and AAV-DJ-DgT at an Ad:AAV ratio of 1:50.

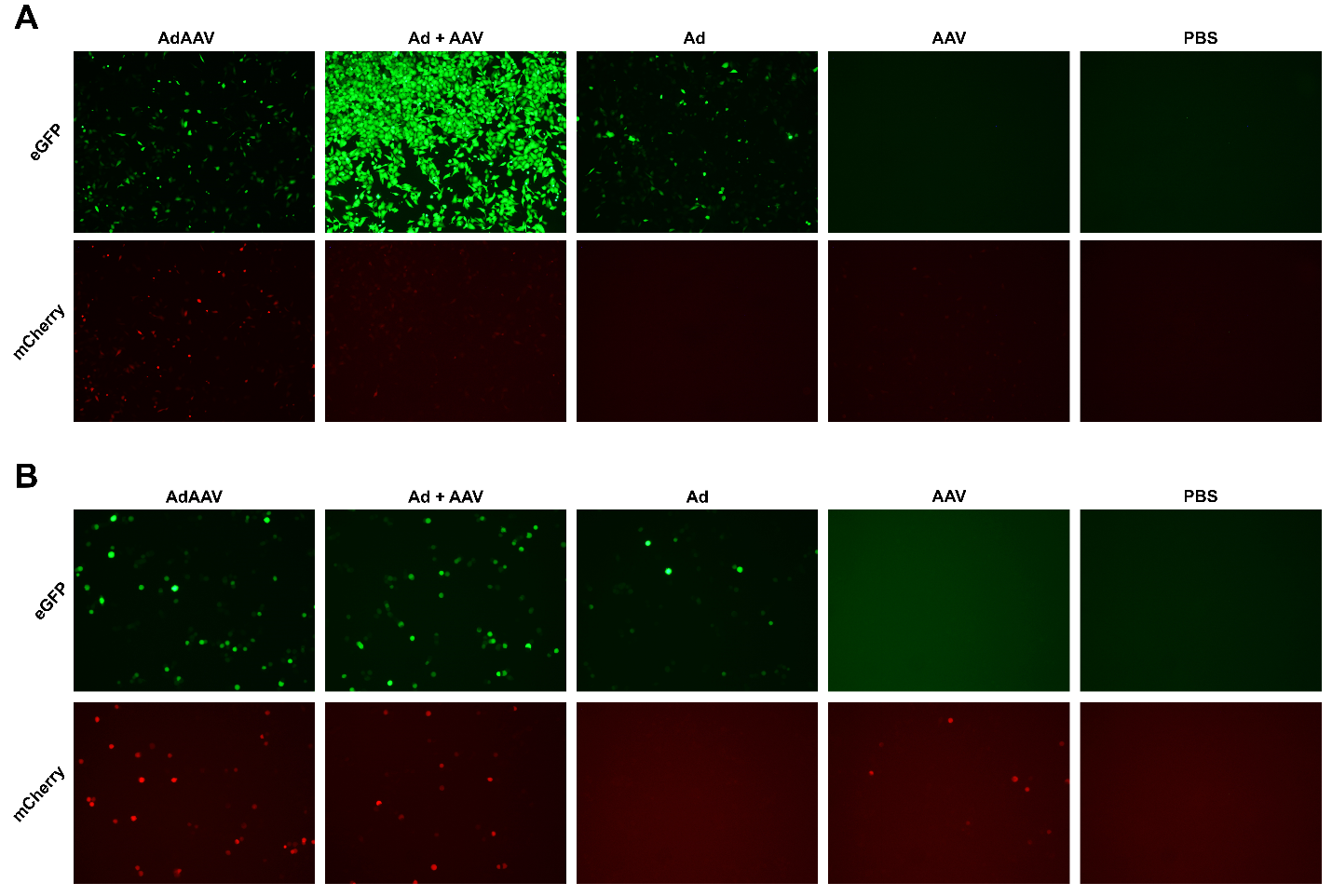

**Supplemental Figure 3 (A)** Fluorescence microscopy images of A549 cells treated with AdAAV, Ad + AAV, Ad alone, AAV alone, and PBS at 3 days post-infection (1:10 Ad to AAV ratio group). Multiplicity of infection (MOI) for the Ad component was 2500 and MOI for the AAV component was 25000, with the exceptions of groups where one of the viruses was excluded. **(B)** Fluorescence microscopy images of A549 cells treated with AdAAV, Ad + AAV, Ad alone, AAV alone, and PBS at 3 days post-infection (1:10 Ad to AAV ratio group). MOI for the Ad component was 500 and MOI for the AAV component was 5000, with the exceptions of groups where one of the viruses was excluded and the PBS groups.

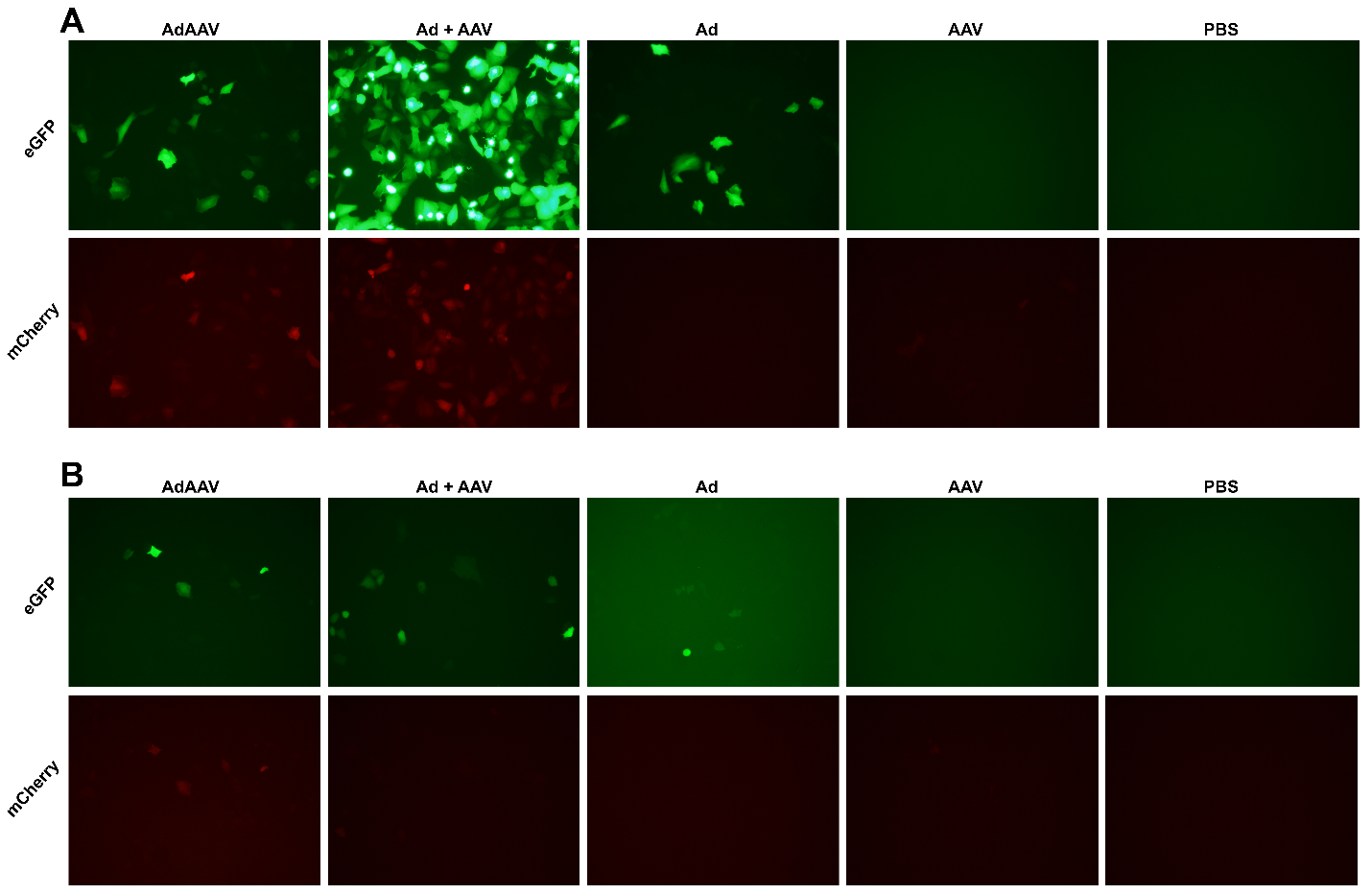

**Supplemental Figure 4 (A)** Fluorescence microscopy images of A549 cells treated with AdAAV, Ad + AAV, Ad alone, AAV alone, and PBS at 3 days post-infection (1:50 Ad to AAV ratio group). Multiplicity of infection (MOI) for the Ad component was 500 and MOI for the AAV component was 25000, with the exceptions of groups where one of the viruses was excluded. **(B)** Fluorescence microscopy images of A549 cells treated with AdAAV, Ad + AAV, Ad alone, AAV alone, and PBS at 3 days post-infection (1:50 Ad to AAV ratio group). MOI for the Ad component was 100 and MOI for the AAV component was 5000, with the exceptions of groups where one of the viruses was excluded and the PBS groups.

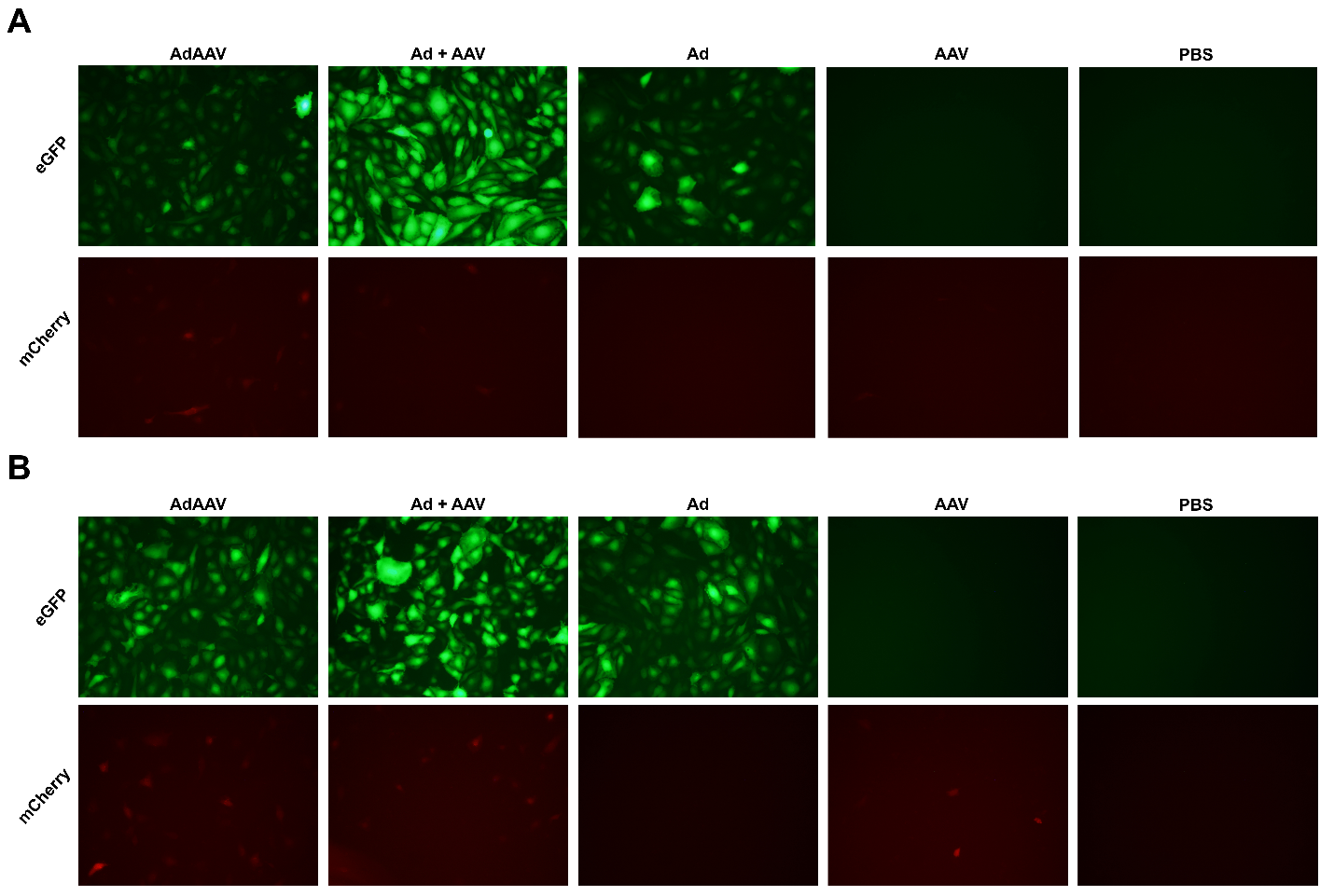

**Supplemental Figure 5 (A)** Fluorescence microscopy images of HUVECs treated with AdAAV, Ad + AAV, Ad alone, AAV alone, and PBS at 3 days post-infection (1:50 Ad to AAV ratio group). Multiplicity of infection (MOI) for the Ad component was 400 and MOI for the AAV component was 20000, with the exceptions of groups where one of the viruses was excluded and the PBS group. **(B)** Fluorescence microscopy images of HUVECs treated with AdAAV, Ad + AAV, Ad alone, AAV alone, and PBS at 3 days post-infection (1:10 Ad to AAV ratio group). Multiplicity of infection (MOI) for the Ad component was 500 and MOI for the AAV component was 5000, with the exceptions of groups where one of the viruses was excluded and the PBS group.

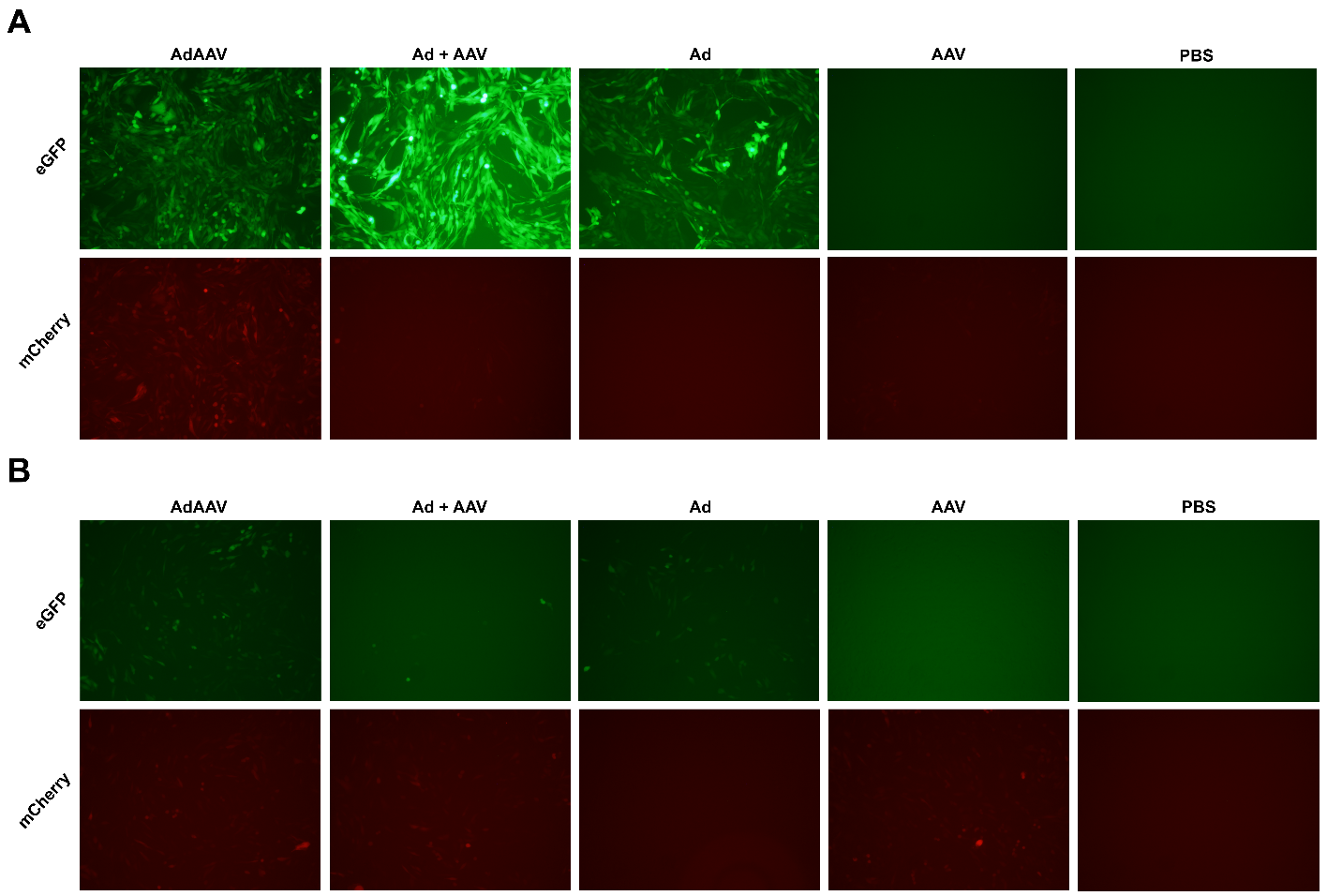

**Supplemental Figure 6 (A)** Fluorescence microscopy images of CHO-CAR cells treated with AdAAV, Ad + AAV, Ad alone, AAV alone, and PBS at 3 days post-infection (1:10 Ad to AAV ratio). Multiplicity of infection (MOI) for the Ad component was 1000 and MOI for the AAV component was 10000, with the exceptions of groups where one of the viruses was excluded and the PBS group. **(B)** Fluorescence microscopy images of CHO cells treated with AdAAV, Ad + AAV, Ad alone, AAV alone, and PBS at 3 days post-infection (1:10 Ad to AAV ratio). Multiplicity of infection (MOI) for the Ad component was 1000 and MOI for the AAV component was 10000, with the exceptions of groups where one of the viruses was excluded and the PBS group.
